# Computational study on ratio-sensing in yeast galactose utilization pathway

**DOI:** 10.1101/2020.05.19.103903

**Authors:** Jiayin Hong, Bo Hua, Michael Springer, Chao Tang

## Abstract

Metabolic networks undergo gene expression regulation in response to external nutrient signals. In microbes, the synthesis of enzymes that are used to transport and catabolize less preferred carbon sources is repressed in the presence of a preferred carbon source. For most microbes, glucose is a preferred carbon source, and it has long been believed that as long as glucose is present in the environment, the expression of genes related to the metabolism of alternative carbon sources is shut down, due to catabolite repression. However, recent studies have shown that the induction of the galactose (GAL) metabolic network does not solely depend on the exhaustion of glucose. Instead, the GAL genes respond to the external concentration ratio of galactose to glucose, a phenomenon of unknown mechanism that we termed ratio-sensing. Using mathematical modeling, we found that ratio-sensing is a general phenomenon that can arise from competition between two carbon sources for shared transporters, between transcription factors for binding to communal regulatory sequences of the target genes, or a combination of the aforementioned two levels of competition. We analyzed how the parameters describing the competitive interaction influenced ratio-sensing behaviors in each scenario and found that the concatenation of both layers of signal integration can expand the dynamical range of ratio-sensing. Finally, we investigated the influence of circuit topology on ratio-sensing and found that incorporating negative auto-regulation and/or coherent feedforward loop motifs to the basic signal integration unit can tune the sensitivity of the response to the external nutrient signals. Our study not only deepened our understanding of how ratio-sensing is achieved in yeast GAL metabolic regulation, but also elucidated design principles for ratio-sensing signal processing that can be used in other biological settings, such as being introduced into circuit designs for synthetic biology applications.

**Author summary:** Microbes make sophisticated choices about the uptake and metabolism of nutrients depending on the variety of nutrient choices available to them in their environment. In the well-studied yeast galactose utilization network, a recent study has shown that galactose metabolic genes respond to the external concentration ratio of galactose to glucose. Using computational models, we showed that this type of phenomenon could arise from a competition between galactose and glucose for transporters, a competition between transcription factors for promoters, or a combination of these two mechanisms. We further revealed the controlling parameters that determined the system sensitivity towards competing input signals and that determined the concentration ratio required to induce the metabolic network in each scenario. Combining competition inhibition at both the transporter level and the transcriptional level can enlarge the ratio-sensing regime, resulting a robust signal integration module. We suspect that modules of this kind may be common in many areas of biology.

## Introduction

Carbon catabolite repression (CCR) is a conserved phenomenon in microorganisms[1–6]. In 1942, Jacques Monod found that when cultured in two carbon sources, bacteria exhibit diauxic growth[7], i.e. they consume the preferred carbon source until it is exhausted, then switch to the less preferred source. This phenomenon was later also observed in yeast[8], for example in the galactose utilization pathway (GAL), which is activated when glucose in the culture is exhausted[9–11]. A recent study has shown that the induction of the GAL pathway in yeast cells is determined not by the absolute level of glucose, but by the concentration ratio of external galactose to external glucose[12]. This mode of induction is termed ratio-sensing. Another study has shown that ratio-sensing is closely related to optimal allocation of protein resources within a cell[13]. A similar ratiometric response, functioning to integrate competing signals, has been identified in the mammalian BMP signaling pathway[14]. Since ratio-sensing responses may have broad importance in biology, we set out to determine what types of general mechanisms can lead to a ratio-sensing response. We constructed simplified mathematical models of competitive binding at both the transporter and transcriptional levels for a simplified version of the GAL pathway in yeast (diagrammed in Fig. 1a), and found that either could be responsible for a ratio-metric response.

**Figure 1:**
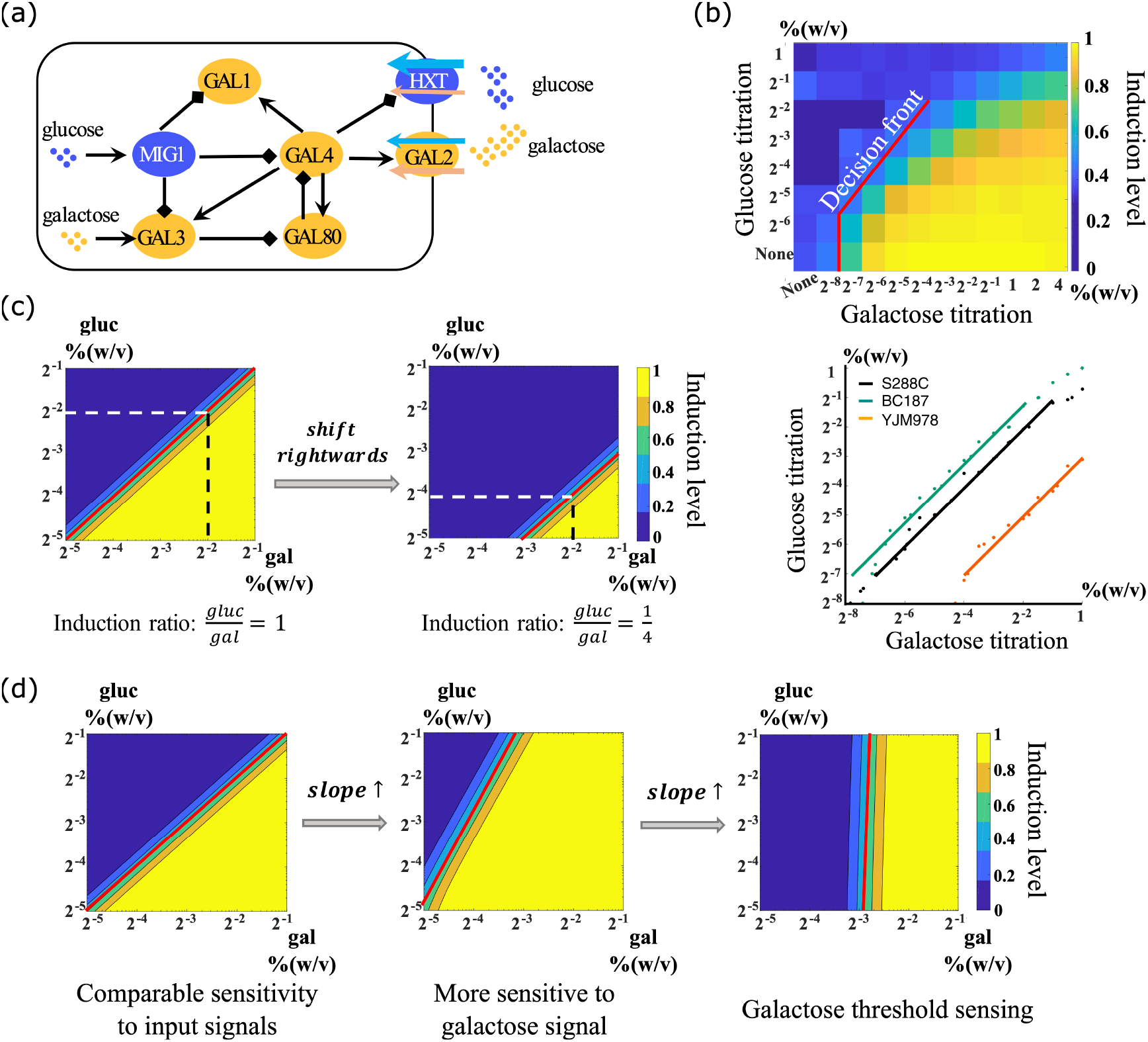
Ratiometric response and the decision front to induce the GAL network. (a) The galactose metabolic gene regulatory network in yeast. (b) Galactose metabolic genes respond to the ratio of external concentrations of galactose and glucose. Different combinations of nutrient concentrations were given to the yeast as indicated by the x and y axes. Top: the color indicates the induction level of the GAL network in each combination of the concentrations. The decision front was defined as the contour line of a fixed induction level. Bottom: three different yeast strains showing three parallel decision fronts. (Data from Ref. [12]). (c) The intercept on the galactose titration axis indicates the induction ratio of glucose to galactose that was required to induce the GAL network. A parallel shift towards higher galactose concentration represents strains that are more susceptible to glucose repression. (d) The slope of the decision front reflects the network relative sensitivity to each nutrient. The contour lines represent specific induction levels of the GAL network. Increasing the slope of the decision front corresponds to elevated sensitivity to the activating galactose signal, and a vertical decision front corresponds to galactose threshold sensing.

To quantitatively determine how glucose depresses GAL pathway induction, we defined a parameter, the “decision condition” that describes the nutrient conditions in which the yeast cells showed half maximum induction of the GAL network. When the decision conditions were plotted on a log-log scale plot of glucose versus galactose concentrations, the curve described by these points was termed the decision front (Fig.1b, upper panel). The slope of the decision front represents the relative sensitivity of the signal integration unit to the competing input signals. When the decision front undergoes a parallel shift, this indicates a change of the concentration ratio that is required to induce the GAL network.

Sugar uptake in yeast is mediated by a variety of transporters. The hexose transporter family (HXT) consists of 17 members, HXT1-HXT17, with varied binding affinity to glucose and other hexoses[15–18]. Gal2p is a galactose permease located in the cellular membrane, which has comparable binding affinity for both galactose and glucose[19]. This suggests that competition at the transporter level between glucose and galactose is possible. Another possible mechanism involves Gal3p, the internal sensor of galactose (see Fig. 1a). When Gal3p binds to galactose it relieves the sequestration of Gal4p by Gal80p, causing transcriptional activation of the downstream GAL metabolic genes including GAL1, GAL2, GAL3, and GAL80[20–26]. If glucose is also present, the internal sensor of glucose, Mig1, transcriptionally represses GAL metabolic genes including GAL1, GAL3 and GAL4[27–30]. In other words, intracellular glucose represses the induction of the GAL pathway through transcriptional inhibition by activating Mig1, indicating that competitive inhibition may also operate at the transcriptional level.

As shown in the diagram (Fig. 1a), there are thus two ways that galactose can be prevented from activating the GAL pathway: high levels of glucose can prevent galactose from entering the cell through competitive binding to the communal transporters HXT and Gal2p, and/or the activation of Mig1 can interfere with transcriptional activation of the pathway by Gal4p. We set out to determine which of these mechanisms can explain the ratio-sensing behavior that was observed experimentally. We found that each of these mechanisms can produce ratio-sensing behavior, and combining both mechanisms produced a robust signal integration mechanism that delivered ratio-sensing behavior over a wide range of input parameters.

## Results

### Decision front

The induction of the galactose metabolic genes in yeast is controlled by both the concentration of galactose and the concentration of glucose in the environment. All lab yeasts and natural yeast isolates studied so far show a ratio-sensing response, but the nutrient conditions required to induce the GAL network vary from one strain to another[12] (Fig.1b, lower panel). To quantitatively study signal integration in the GAL network, we built coarse-grained ODE models (Methods and Models) that described the reactions that occur at the transporter level and the transcriptional level, and simulated the behavior of the network when induced by double gradients of glucose and galactose (Fig.1c, 1d). The contour lines in the double titration graph delineate specific induction levels of the GAL network. Each point on a contour line represents a combination of glucose and galactose concentrations that results in the same induction level of the GAL pathway. We call this contour line the decision front. To better understand the possible types of behaviors from our model, we will describe what several different types of qualitative changes would mean in terms of glucose and galactose regulation. This explanation will help to understand the behaviors observed throughout the paper.

Each yeast isolate has a characteristic decision front, representing the concentration ratio required to induce the GAL network in that strain (Fig. 1c). As the decision front shifts rightwards along the galactose titration axis, the induction ratio of glucose to galactose decreases from 1 to ¼, indicating that the strains require more galactose to be present in their environment before they induce the GAL network.

The slope of the decision front reflects how sensitive the yeast strain is to the presence of galactose and glucose (Fig.1d). An increased slope of the decision front reflects greater sensitivity to the activating galactose signal than to the suppressing glucose signal. Strains with decision front almost vertical to the galactose titration axis indicate that induction of the GAL pathway is almost independent of external glucose in these strains. In contrast, a decreased slope of the decision front indicates increased sensitivity to suppression by glucose.

### Realizing ratio-sensing by competitive binding to the communal transporter

How is ratio-sensing implemented at the molecular level? We first focused on the possibility that transporter competition was responsible. We abstracted the uptake of glucose and galactose through HXT and Gal2p as a simplified mathematical model (see Methods and Models for details). Galactose and glucose are transported into the cell by a shared transporter, with binding coefficients of *K*_*gal*_ and *K*_*gluc*_, and cooperativity coefficients of *n*_*gal*_ and *n*_*gluc*_, respectively (Fig.2a). We assumed that the total number of transporters within a cell was constant and had three states: bound by glucose, bound by galactose, or empty. We simulated transport of both sugars under these conditions, and assumed that the GAL network would be induced when the intracellular galactose level was above a threshold concentration of galactose. Solving the equations at steady state yielded an expression describing the intracellular galactose level in terms of external glucose and galactose.

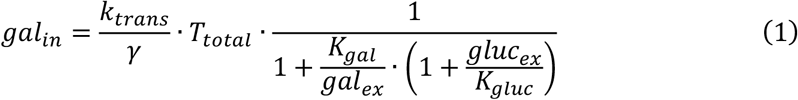

**Figure 2:**
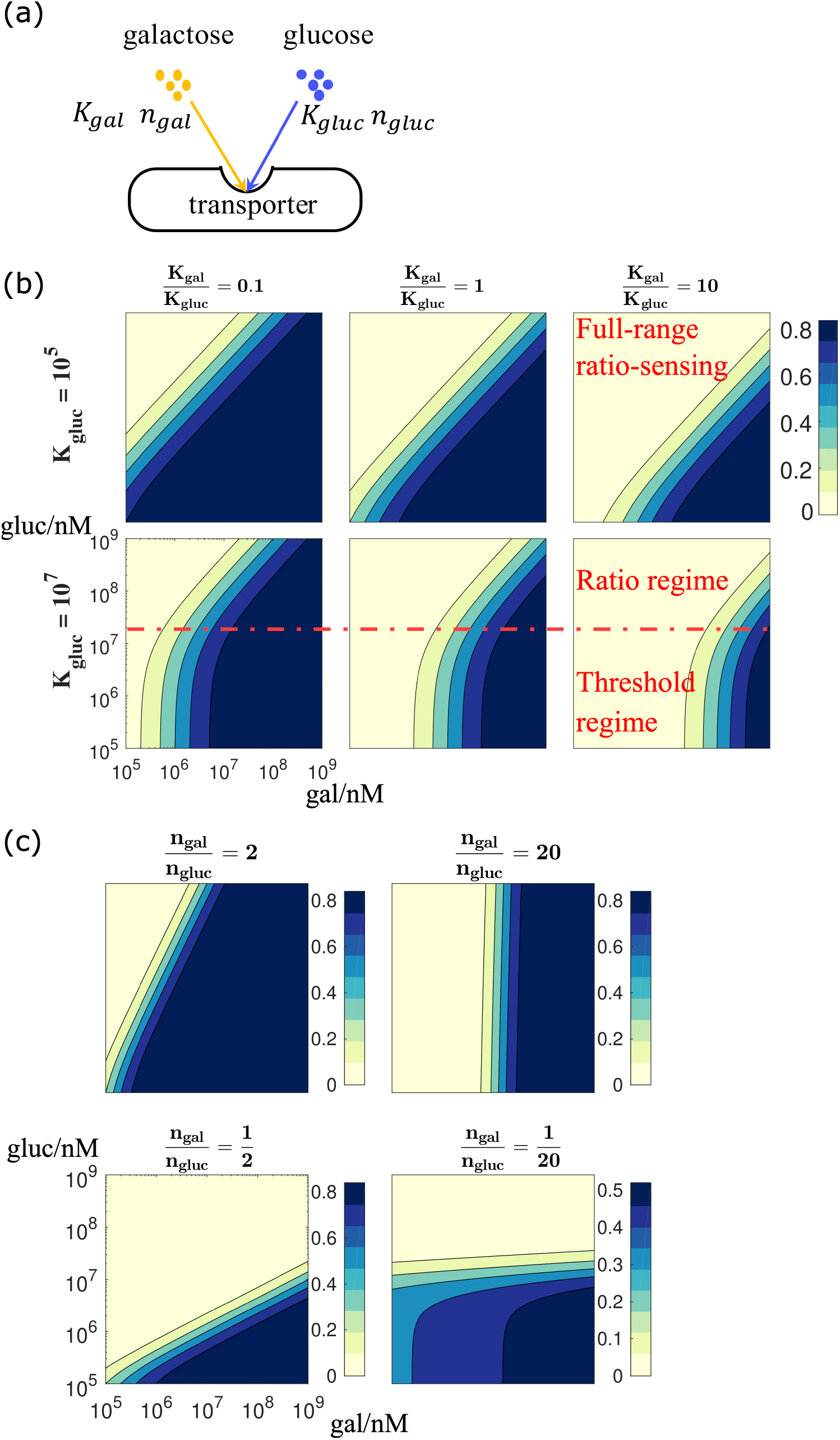
Realizing ratio-sensing by competitive binding to the communal transporter. (a) The transporter level competitive binding model. (b) Simulations of the induction level of the GAL pathway in the transporter level model. While keeping *K*_*gluc*_ unchanged, increasing the relative binding affinity 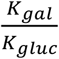 shifts the decision front towards higher galactose concentration. The upper panel parameter regime with *K*_*gluc*_ = 10^5^ exhibits a full-range ratiometric response, whereas the lower panel parameter regime with *K*_*gluc*_ = 10^7^ exhibits a compound signal integration mode, i.e. galactose threshold sensing at poor nutrient condition and ratio-sensing at rich nutrient condition. (c) The network sensitivity to input signals is determined by relative binding cooperativity. Increasing the relative cooperativity makes the signal integration unit more sensitive to activating galactose signal, whereas decreasing the relative cooperativity makes the signal integration unit more sensitive to repressing glucose signal. In the extreme case, where the decision front is almost vertical to the galactose titration axis 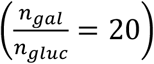, the signal integration unit becomes a galactose threshold sensor. When the decision front is almost parallel to the galactose titration axis 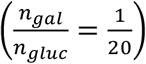, the signal integration unit becomes a glucose threshold sensor.

In equation (1), *k*_*trans*_ is the maximal transportation rate through the given transporter, *γ* is the turn-over rate of the sugar within a cell, *T*_*total*_ is the total number of transporters expressed on the membrane, *K*_*gal*_ and *K*_*gluc*_ are the binding coefficients for galactose and glucose respectively, and *gal*_*ex*_ and *gluc*_*ex*_ are the concentrations of the external carbon sources. Equation (1) suggests that when the maximal transportation rate and turn-over rate within a cell are fixed, a ratio-sensing regime exists as long as the binding affinity coefficient between glucose and the transporter is much smaller than the external glucose concentration (*K*_*gluc*_ ≪ *gluc*_*ex*_), i.e. the external glucose level is high enough to saturate the transporters.

Next, we derived the expression of the decision front in log-log scale as follows:

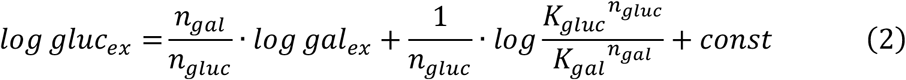

where *n*_*gal*_ and *n*_*gluc*_ are the cooperativity coefficients of galactose and glucose binding to the transporter, respectively. From equation (2) we know that the intercept on the galactose titration axis is determined by the relative binding affinity between the two sugars and the communal transporters. This explains why the decision front shifted to a higher 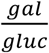 ratio as we increased 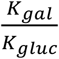 (Fig.2b). This makes intuitive sense: the stronger the binding between glucose and the transporter, or the weaker the binding between galactose and the transporter, the more the GAL network should be inhibited by glucose and the higher the level of external galactose required to induce the GAL pathway.

Note that when the condition *K*_*gluc*_ ≪ *gluc*_*ex*_ is met, we observe a ratio regime across the full physiological plausible range of double sugar titration. In contrast, when *K*_*gluc*_ ≪ *gluc*_*ex*_ is violated, the system exhibits a compound signal integration mode, i.e. at low concentrations of both glucose and galactose, the GAL pathway responds solely to the external galactose signal. In other words, we observe a galactose threshold sensing at low concentrations of sugars, but at high concentrations of both sugars we see a ratio-sensing response. This is consistent with the experimental observations reported by Escalante-Chong et al[12]. The analytical derivation also shows that the slope of the decision front is determined by the relative cooperativity of galactose and glucose binding to the transporter. Varying the value of 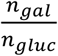 changed the slope of the decision front, and when 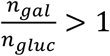, the system is more sensitive to the external galactose signal. With 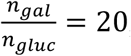, the system becomes almost independent of glucose concentrations, i.e. galactose threshold sensing, for a portion of the phase space as seen in Figure 2c. In contrast, when 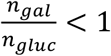, the system is more sensitive to the external glucose signal. With 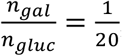, the system becomes almost solely responsive to the variation of glucose concentration, i.e. glucose threshold sensing (Fig. 2c).

### Realizing ratio-sensing by transcriptional inhibition of GAL metabolic genes

Apart from the cross-talk at the transporter level, glucose also inhibits the expression of GAL genes through the activation of the transcription factor Mig1. We generalized this mechanism as a two-step reaction, and explored whether it can also produce a ratio-sensing response (Fig. 3a). In the first step of this reaction system, both the activating signal (galactose) and the inhibiting signal (glucose) bind to their corresponding intracellular sensors, forming the activator (Gal3p*) and the repressor (Mig1*). In the second step, either the activator or the repressor binds to a cis-regulatory element (CRE) of the readout (GAL1), where the conflicting effect was integrated at the transcriptional level. Again, we used the law of mass action to model the association and dissociation between the sugars and the sensors, and that between the sensors and the CRE (see Methods and Models). Here, we assumed that the copy number of the GAL1 gene was constant within a cell, and the CRE of GAL1 had three possible states: bound by the activator and transcriptionally active; bound by the repressor and thus transcriptionally inhibited; and free such that it is not inhibited but not transcribing. Because the biochemical reactions of binding and unbinding between sugars and sensors are much more rapid than the process of transcription[31], we used separation of time-scale in our model and obtained a quasi-equilibrium approximation for the functional forms of the activator and the repressor:

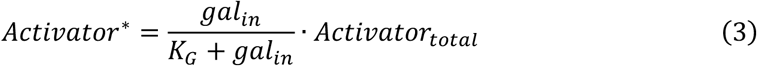

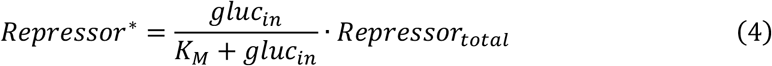

where *K*_*G*_ and *K*_*M*_ are the binding affinity of galactose and glucose to their corresponding intracellular sensors, respectively, and *Activator*_*total*_ and *Repressor*_*total*_ are the total amount (free forms plus functional forms) of the activator and the repressor.

**Figure 3:**
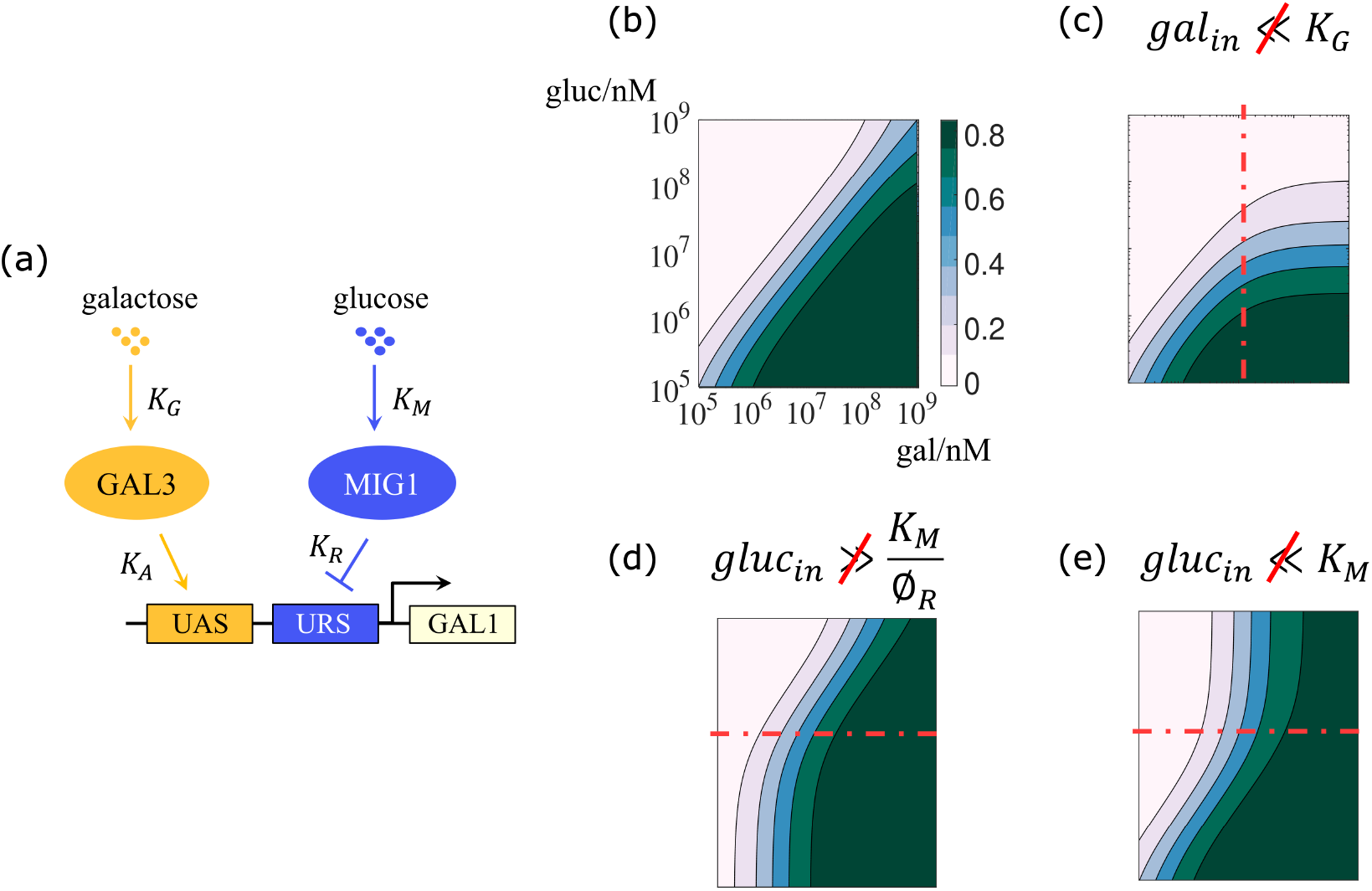
Realizing ratio-sensing by transcriptional inhibition of GAL metabolic genes. (a) The transcriptional level competitive inhibition model. (b) The transcriptional level model can give rise to a full-range ratiometric response in a reasonable parameter regime. (c) When the requirement *gal*_*in*_ ≪ *K*_*G*_ is violated, the signal integration unit exhibits ratio-sensing at low carbon source concentrations, and glucose threshold sensing at high carbon source concentrations. (d) When the requirement 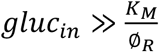 is violated, the signal integration unit exhibits galactose threshold sensing at low carbon source concentrations, and ratio-sensing at high carbon source concentrations. (e) When the requirement *gluc*_*in*_ ≪ *K*_*M*_ is violated, the signal integration unit exhibits ratio-sensing at low carbon source concentrations, and galactose threshold sensing at high carbon source concentrations.

When the association and dissociation between the cis- and trans-regulatory elements of GAL1 reaches equilibrium, the fraction of actively-transcribed genes among the total number of GAL1 copies is:

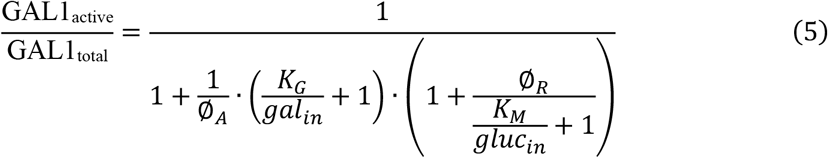

where 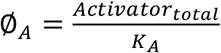 and 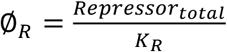. *K*_*A*_ and *K*_*R*_ are the dissociation constants of the activator and the repressor binding to the CRE of GAL1, respectively. Analytical derivation showed that a ratio-sensing regime emerged when the following requirements were simultaneously met: *gal*_*in*_ ≪ *K*_*G*_ and 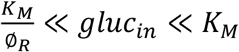. The decision front is described by:

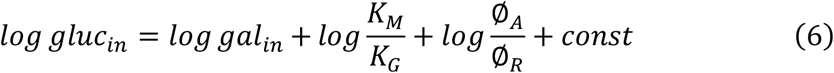

Equation (6) indicates that the slope of the decision front in a basic signal integration circuit is fixed at 1, and the intercept on the galactose titration axis is determined by 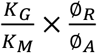, where *K*_*G*_ and *K*_*M*_ represent the binding affinity of galactose to the activator, and of glucose to the repressor, respectively. ∅_*R*_ measures the repression capacity of the repressor, with greater values signaling stronger inhibition. Similarly, ∅_*A*_ represents the activation capacity of the activator.

Numerical simulations of this system shows that ratio-sensing is achieved if *gal*_*in*_ ≪ *K*_*G*_ and 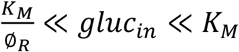 are simultaneously met (Fig. 3b). Violation of any of these requirements results in a combination of ratio-sensing and threshold sensing. Specifically, when *gal*_*in*_ ≪ *K*_*G*_ is violated, the system responds to the ratio of the two nutrient signals at low input concentrations, but only responds to changes in glucose levels at high input concentrations (Fig. 3c). When 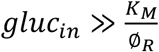 is violated, the system exhibits galactose threshold sensing at low input concentrations, but responds to the concentration ratio at high input concentrations (Fig. 3d). Lastly, when *gluc*_*in*_ ≪ *K*_*M*_ is violated, the system exhibits ratio-sensing at low input concentrations, but responds only to changes in galactose levels at high input concentrations (Fig. 3e).

### A combination of both transporter-level and transcription-level competition expands the dynamical range of ratio-sensing

Our analysis showed that ratio-sensing could arise from either competitive binding to the shared transporters alone, or from transcriptional regulation of the GAL metabolic genes alone. Since glucose and galactose do compete for Gal2p and HXTs, and transcriptional inhibition of the GAL metabolic genes by Mig1 also occurs, we were curious about whether there is a benefit to yeast cells that could use both mechanisms for ratiometric signal integration. We therefore modeled a combination of both mechanisms, using the output of the competitive transporter model as the input of the transcriptional regulation model, to investigate how the combined system behaved.

We combined two typical signal integration behaviors from the transporter model (Fig. 4a) with four typical signal integration behaviors of transcriptional model (Fig. 4b), and simulated the response of the resultant eight combinations (Fig. 4c). We found that, no matter whether the single integration modes were ratio-sensing or a mix of ratio-sensing and threshold sensing, all eight combinations exhibited ratio-sensing response. In other words, combining both mechanisms together expanded the dynamical range over which a ratiometric response was observed.

**Figure 4:**
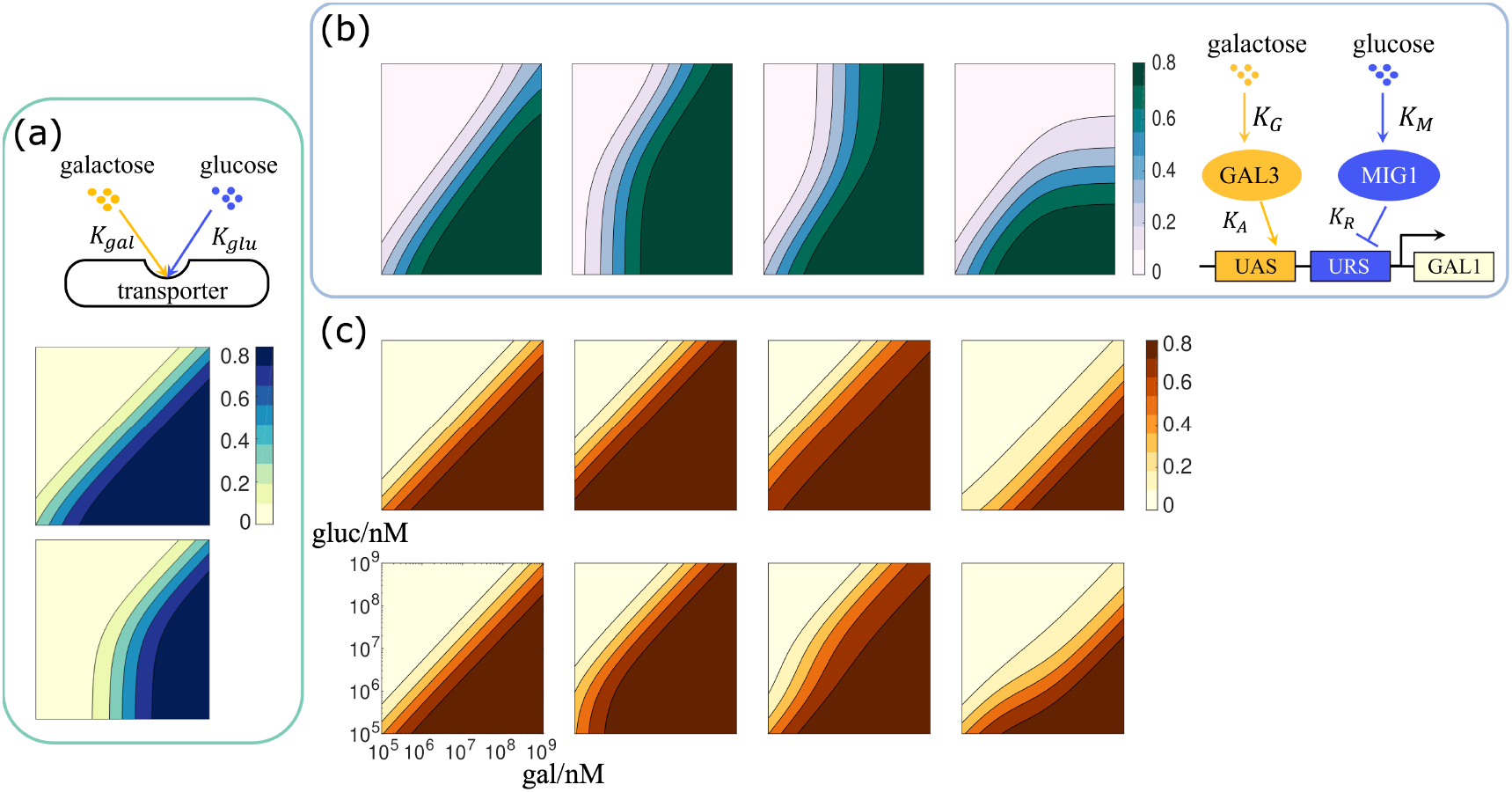
Concatenating the dual layers of signal integration expanded the dynamical range of ratio-sensing. (a) The two typical signal response patterns in the transporter level model. The upper pattern exhibits full-range ratio-sensing while the lower one exhibits galactose threshold sensing combined with ratio-sensing. (b) The four typical signal response patterns in the transcriptional level model specified in Fig. 3(b)-(e). (c) The output signal response patterns of the concatenation model all exhibit full-range ratio-sensing, despite the fact that some response patterns in either the upstream transporter layer or the downstream transcriptional layer showed compound signal integration.

To determine the mechanism underlying this effect, we analyzed the requirements for ratio-sensing in the combined model. The transcriptional regulation model alone generates a ratiometric response only when *gal*_*in*_ ≪ *K*_*G*_ and 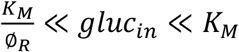. In the combined model, both *gal*_*in*_ and *gluc*_*in*_ are determined by the behavior of the hexose transporters. Hence, we used equation (1) and

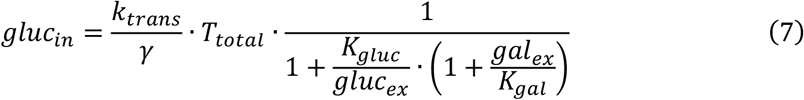

to substitute *gal*_*in*_ and *gluc*_*in*_, yielding the following requirements for ratio-sensing in the concatenation model:

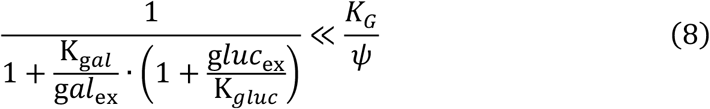

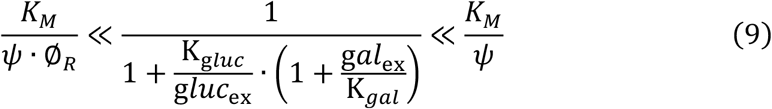

where 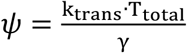 represents the sugar transportation capacity.

Note that 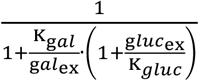 and 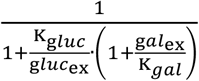 are always smaller than 1, which means that as long as *K*_*G*_ ≫ *ψ* and *K*_*M*_ ≫ *ψ* are satisfied and ∅_*R*_ is large, the system will respond in a ratiometric manner.

This analysis implies that when the external glucose concentration is low, causing the upstream transporter circuit to fall into a galactose threshold sensing pattern, the system is nevertheless able to realize ratio-sensing if the input sugars bind to their corresponding internal sensors with low affinity, so that *K*_*G*_ ≫ *ψ* and *K*_*M*_ ≫ *ψ* are satisfied. Conversely, when the adjustable capacity of the downstream transcriptional regulatory layer is limited, causing the CRE of GAL1 to be saturated by either the activator or the repressor, the system can realize ratio-sensing by reducing the production of transporters, so that *K*_*G*_ ≫ *ψ* and *K*_*M*_ ≫ *ψ* are again satisfied. Taken together, the combination of the two layers of molecular circuits increases the flexibility of signal integration, giving rise to an expanded ratio-sensing regime.

### Incorporating network motifs to the basic signal integration unit altered the network sensitivity to input signals

In the model of competitive binding at the transporter level, we showed that the relative binding cooperativity of the competing sugars determined the sensitivity of the signal integration circuit to external nutrient signals, that is, the slope of the decision front can change as the system becomes more sensitive to one sugar than the other (Fig. 2c). This stands in contrast to the transcription-level model, which always has a decision front with slope approximately equal to 1, i.e. the simple signal integration unit we modeled had comparable sensitivity to both of the competing input signals (Fig. 3b). We were curious whether it is possible to alter the behavior of the system by changing the topology of the signal integration unit.

There are recurring regulatory patterns that appear in transcriptional networks much more frequently than would be expected for random networks. These patterns, called network motifs, have been found in a range of organisms from microbes to mammalian cells[32–34]. We incorporated selected network motifs into our original model and tested how this changed the response to the nutrient stimuli. While the slope of the decision front in our original model was approximately 1 (Fig.5a), we found that introducing negative autoregulation[35,36] at the activator node, and/or introducing a negative regulatory edge from the repressor to the activator, forming a coherent feedforward loop[37], decreased the slope of the decision front, and made the signal integration unit more sensitive to the repressing nutrient signal (Fig.5b). In contrast, introducing negative autoregulation at the repressor node, and/or introducing a negative regulatory edge from the activator to the repressor, forming a coherent feedforward loop, increased the slope of the decision front, and made the signal integration unit more sensitive to the activating nutrient signal (Fig.5c). When we analyzed the steady state induction level of the pathway, we found that these network motifs not only altered the sensitivity to the input signals, but also changed the robustness of the response when intracellular protein levels were allowed to fluctuate (see Methods and Models for details).

**Figure 5:**
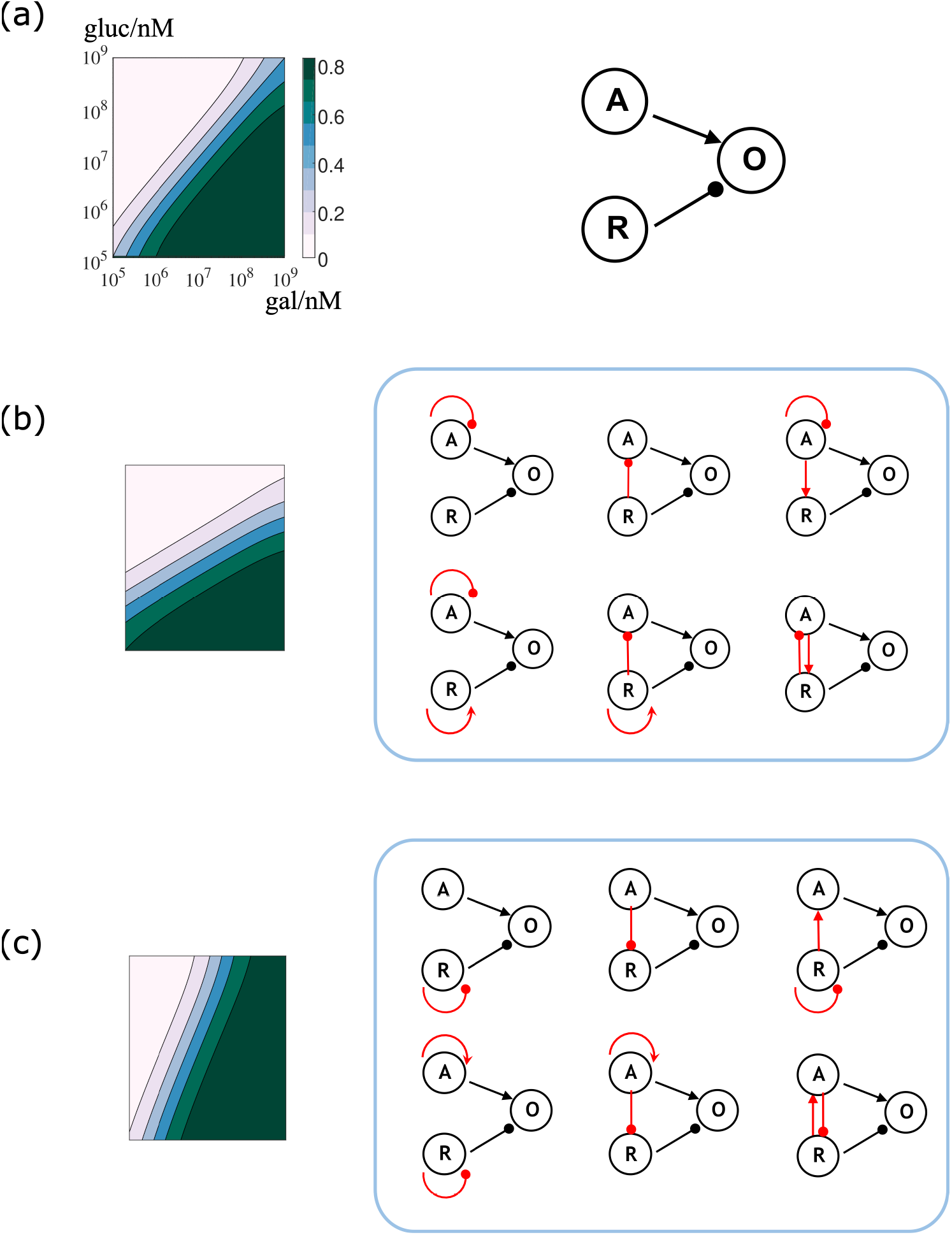
Incorporating network motifs to the basic signal integration unit altered the network sensitivity to input signals. (a) The basic signal integration unit consists of the activator (node A), the repressor (node R), and the output gene (node O). The slope of the decision front for this network is approximately equal to 1. (b) Network topologies that halve the slope of the decision front, i.e. are more susceptible to inhibition by glucose. These topologies involved negative auto-regulation at the activator node and/or an inhibiting edge from the repressor to the activator. (c) Network topologies that double the slope of the decision front, i.e. are more sensitive to activation by galactose. These topologies involve negative auto-regulation at the repressor node and/or an inhibiting edge from the activator to the repressor.

## Discussion

Carbon catabolite repression is a conserved phenomenon across microorganisms, and it has long been established that the yeast GAL pathway is induced when the external glucose concentration drops. Recent studies have shown that yeast cells respond not to a simple glucose threshold, as previously believed, but to the ratio of the external concentrations of galactose and glucose[12]. Our research aimed to determine which molecular circuits could be responsible for ratiometric sensing of nutrient signals. We found that two segments of the sugar uptake/response pathway each had features that could give rise to a ratiometric response. The first of these is the family of transporters that take up the two sugars. Nutrient transporters are not generally considered to be part of signaling pathways or responsible for signal processing, but we found that competition between sugars for transporter binding could explain the ratio-sensing response. The relative binding affinity of the competing carbon sources for the communal transporter determined the concentration ratio for the induction of the GAL pathway. If the two sugars are each allowed to show cooperativity in binding to the transporter, the relative degree of cooperativity determined which sugar the pathway responded to more sensitively. When the cooperativity of galactose binding was overwhelmingly greater than that of glucose binding, the pathway behaved as a galactose threshold sensor; conversely, when the cooperativity of glucose binding was much greater than that of galactose, the pathway only responded to variations in glucose concentration.

The second segment of the sugar uptake/response pathway that can give a ratiometric response is the regulation of transcription. This molecular circuit consists of two sequential reactions, the binding of the intracellular nutrients to their internal sensors, producing an activator complex and a repressor complex, and the binding of these activator/repressor complexes to the cis-regulatory element (CRE) of the GAL1 gene. In this circuit many factors are important for the induction ratio: the relative binding strength between the competing sugars and their corresponding intracellular sensors, the total amount of each type of sensor, and the relative binding strength of the activator complex and the repressor complex to the GAL1 CRE. The behavior of this circuit can be further modulated by introducing negative auto-regulation and/or coherent feedforward loop motifs.

Since both of these elements are found together in the GAL control pathway, we examined how they would work when combined. Concatenating the transporter layer and the transcriptional regulatory layer resulted in ratio-sensing behavior with an expanded dynamical range and increased flexibility. This double-layer model can accommodate a wide and physiologically plausible regime for ratio-sensing by adjusting either the transportation capacity of the transporters or the binding strength between the sugars and their corresponding intracellular sensors. Different strains of yeast that have encountered different conditions during their evolution would therefore be expected to show variation in these parameters.

Our research reveals simple design principles for ratio-sensing signal processing which may be helpful in identifying such behavior in other systems, and can be used to design systems with desirable properties for synthetic biology applications.

## Methods and Models

### Modeling competitive binding at transporter level

We assumed that the total amount of membrane transporters was a constant, and that the transporters could be free, bound by galactose, or bound by glucose. Thus:

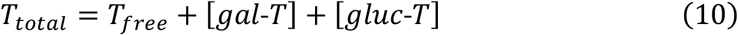

Using the law of mass action, the levels of intracellular galactose, intracellular glucose, transporters bound by galactose, transporters bound by glucose, and free transporters are related as follows:

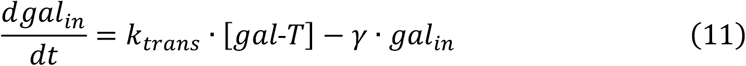

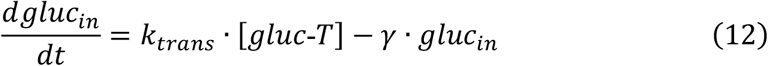

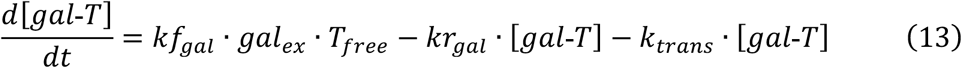

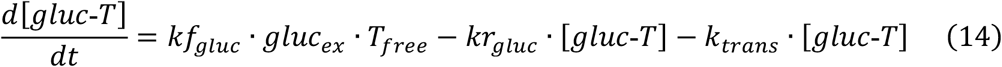

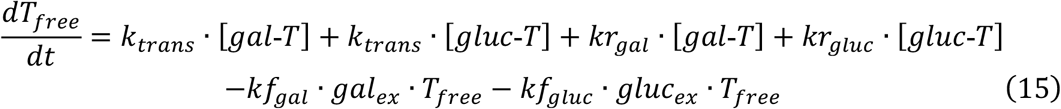

When the above reactions reach steady state, the left sides of equations (11) to (15) are all equal to 0, and we can derive the intracellular galactose level and intracellular glucose level at steady state. This calculation produced equation (1) and equation (7) in the main text.

To derive the expression of decision front and solve for the slope and the intercept on *gal* titration axis, we set the intracellular galactose level equal to a constant (*const*):

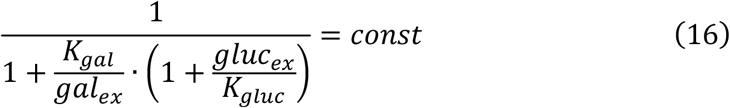

When *gluc*_*ex*_ ≫ *K*_*gluc*_, equation (16) in log-log scale is approximately:

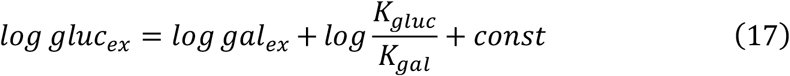

A generalized form of equation (17) incorporating cooperativity was shown in equation (2).

### Modeling competitive inhibition at transcriptional level

We assumed that GAL1 gene copies within a cell are constant, and that the cis-regulatory element of GAL1 could be bound by the activator, bound by the repressor, or free. So, we have,

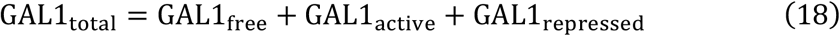

When the association and dissociation between regulatory factors and CRE reach equilibrium, we have,

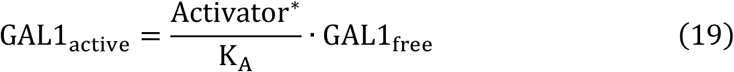

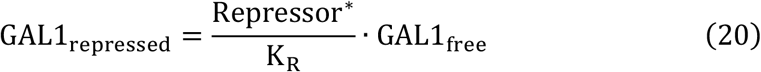

where 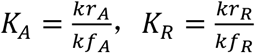.

Combining equation (3), equation (4), and the above equations, we could derive equation (5), which describes the actively transcribed fraction of GAL1 gene copies, as well as equation (6), which gives the formula for decision front in the transcriptional level model.

### Introducing network motifs to transcriptional level model

The generalized expression for the actively transcribed fraction of given gene copies is,

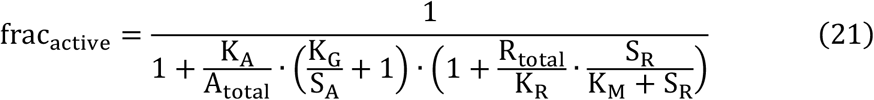

where S_A_ and S_R_ represent the concentration of activating input signal and of repressing input signal, respectively.

We first considered introducing negative auto-regulation to the activator node, which means that making more activators would lead to inhibition of activator expression. We used the following equation to model negative auto-regulation at the activator node:

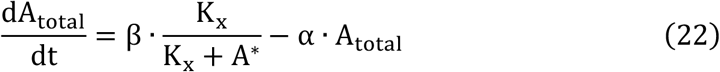

where A* was the functional form of the activator upon binding by S_A_, and K_t_ was the concentration of A* when the synthesis of the activator reached half-maximum inhibition, i.e. the synthesis rate of the activator was half of the maximum synthesis rate. At steady state, we find

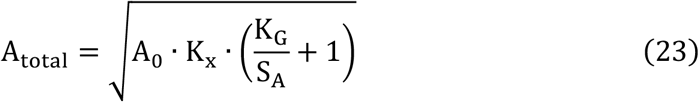

where 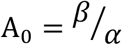, the expression level of the activator without introducing negative auto-regulation. Within the ratio-sensing regime, substituting equation (23) to equation (21) yields

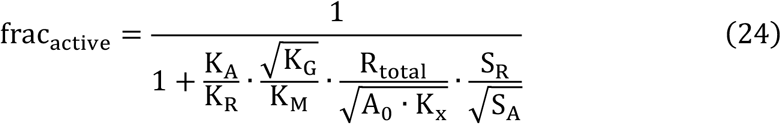

Meanwhile, we could also solve the formula for the decision front as follows:

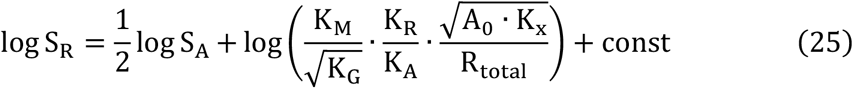

Equation (25) indicates that incorporating negative auto-regulation at the activator node would halve the slope of the decision front in log-log scale, in other words, the network would become more sensitive to the inhibiting glucose signal. The intercept on the galactose titration axis also becomes dependent on the square root of *K*_*G*_ and A_0_, which means that the decision to induce the network is more robust to variation of the binding affinity between galactose and the activator, as well as more robust to the fluctuation of the activator level.

Next, we considered the introduction of positive auto-regulation to the activator node. This gives

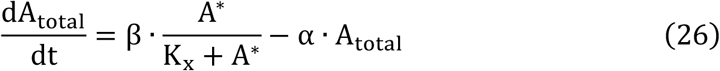

At steady state,

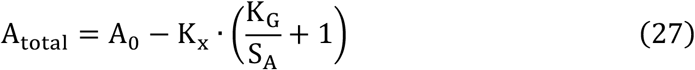

Within the ratio-sensing regime, substituting equation (27) to equation (21) yields

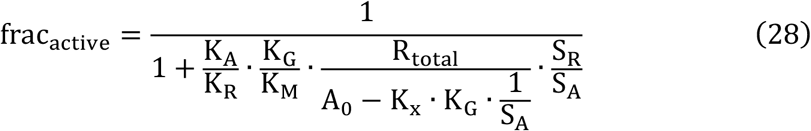

And the formula for the decision front is

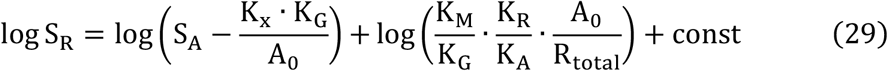

Equation (29) suggests that incorporating positive auto-regulation to the activator changes neither the slope of the decision front, nor the sensitivity to input signals.

When negative auto-regulation of the repressor is introduced,

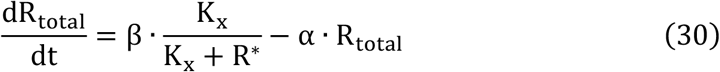

where R* represents the functional form of the repressor upon binding by glucose. Again, we solved for total repressor when the system reaches steady state:

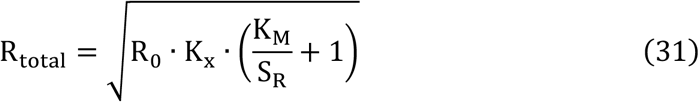

where 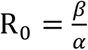, the expression level of the repressor without the negative auto-regulation motif. Within the ratio-sensing regime, substituting equation (31) to equation (21) yields

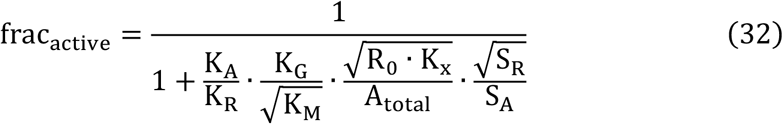

The formula for the decision front is

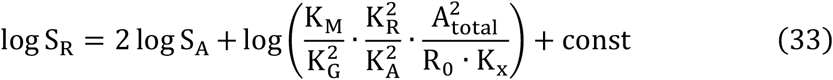

Equation (33) implies that incorporating negative auto-regulation at the repressor node doubles the slope of the decision front, which means that the network becomes more sensitive to the activating galactose signal. Meanwhile, the intercept on the galactose titration axis becomes dependent on the square of *K*_*G*_, *K*_*R*_, *K*_*A*_ and A_*total*_, so that the decision to induce the network is more susceptible to variations in the binding affinity between galactose and the activator, or the binding affinity between the regulator and the cis-regulatory element of GAL1, as well as more sensitive to fluctuations in the activator level.

Similar to the activator node, introducing positive auto-regulation to the repressor node does not change the slope of the decision front. The actively transcribed gene fraction and the formula for the decision front are given by:

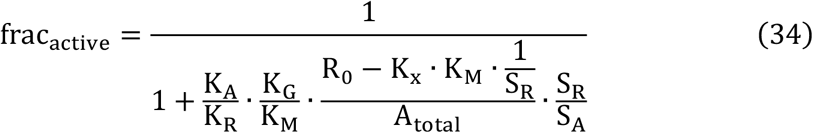

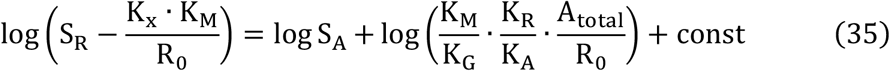

We next sought to understand how feedforward loops change the integration of the different signals. First, consider the case where the activator promotes the expression of the repressor, forming a type I incoherent feedforward loop (IFFL type I). We have

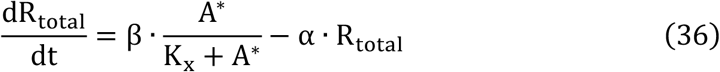

When cells reach steady state:

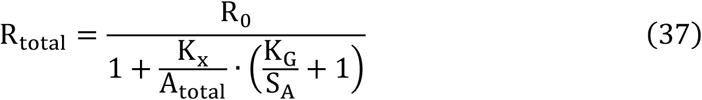

Within the ratio-sensing regime, substituting equation (37) to equation (21) yields

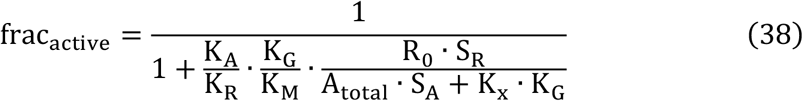

Solving the formula for the decision front:

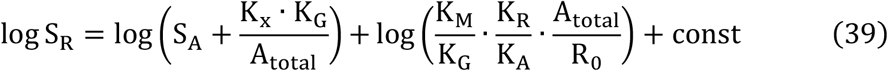

We found that introducing IFFL type I did not change the network sensitivity to input signals.

We next considered the case where the activator inhibited the expression of the repressor, forming type I coherent feedforward loop (CFFL type I). We have

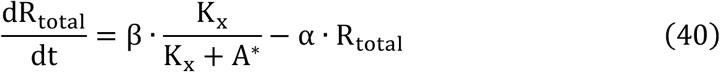

And the total amount of the repressor at steady state:

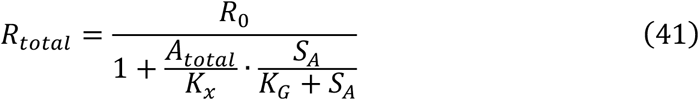

Within the ratio-sensing regime, substituting equation (41) to equation (21) yields

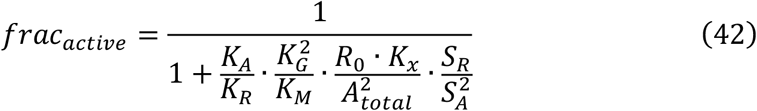

And the formula for the decision front:

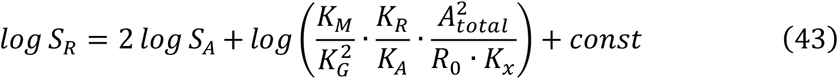

Equation (43) indicates that incorporating CFFL type I doubles the slope of the decision front in log-log scale, in other words, the network becomes more sensitive to the activating galactose signal. Meanwhile, the intercept on the galactose titration axis becomes dependent on the square of *K*_*G*_ and A_total_, which means the decision to induce the network is more susceptible to variations in the binding affinity between galactose and the activator, as well as more sensitive to fluctuations in the activator level.

We then introduced an activating edge from the repressor to the activator, forming a type II incoherent feedforward loop (IFFL type II). The dynamical differential equation and steady state level of the activator are:

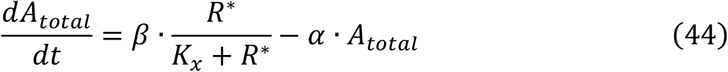

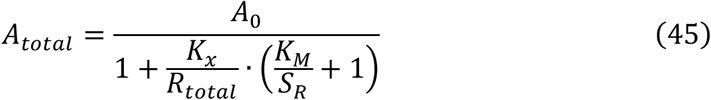

Within the ratio-sensing regime, substituting equation (45) to equation (21) yields

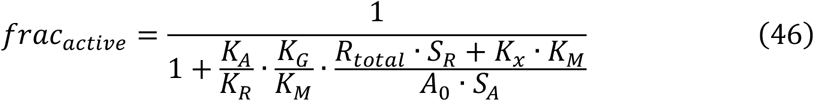

And the formula of the decision front:

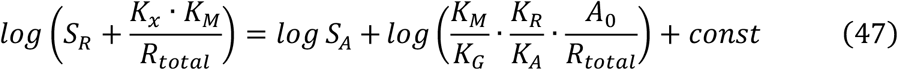

Equation (47) indicates that incorporating IFFL type II does not change the network sensitivity to input signals.

Finally, we introduced an inhibitory edge from the repressor to the activator, forming a type II coherent feedforward loop (CFFL type II). The dynamical differential equation and steady state level of the activator are:

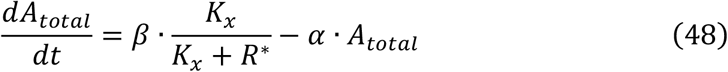

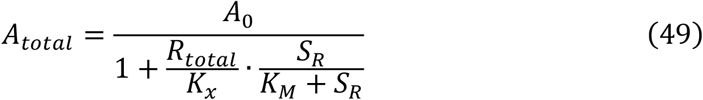

Within the ratio-sensing regime, substituting equation (49) to equation (21) yields

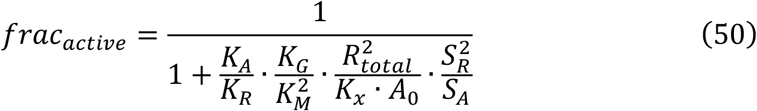

And the formula of the decision front:

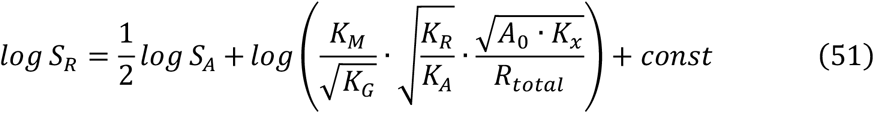

Equation (51) implies that incorporating CFFL type II halves the slope of the decision front in log-log scale, in other words, the network is more sensitive to the inhibiting glucose signal. The intercept on the galactose titration axis is dependent on the square roots of *K*_*G*_, *K*_*R*_, *K*_*A*_ and A_0_, which means that the decision to induce the network is more robust to variations in the binding affinity between galactose and the activator, and between the regulators and the cis-regulatory element, as well as more robust to the fluctuation of the activator level.

We further combined auto-regulation with feedforward loops, and studied how they affected network sensitivity to input signals. For either the activator or the repressor, there could be five regulatory states in total: positive auto-regulation, negative auto-regulation, coherent feedforward loop, incoherent feedforward loop, and without any regulation. As the regulatory states between the activator and the repressor were independent, there were twenty-five possible configurations for the downstream gene expression output. Among them, we found that six configurations that halved the slope of the decision front, i.e. made the system more sensitive to inhibiting glucose signals. These six configurations involved negative auto-regulation of the activator, and/or inhibitory edges from the repressor to the activator. Similarly, there were six configurations that doubled the slope of the decision front, i.e. made the system more sensitive to activating galactose signals. These six configurations involved negative auto-regulation of the repressor, and/or inhibitory edges from the activator to the repressor.

## Acknowledgments

We thank Xiaojing Yang and Yihan Lin for helpful discussions and comments, and Becky Ward for critically reading the manuscript.

